# Arf GTPase-Activating proteins ADAP1 and ARAP1 regulate incorporation of CD63 in multivesicular bodies

**DOI:** 10.1101/2024.02.15.580439

**Authors:** Kasumi Suzuki, Yoshitaka Okawa, Sharmin Akter, Haruki Ito, Yoko Shiba

## Abstract

Arf GTPase-Activating proteins (ArfGAPs) mediate the hydrolysis of GTP bound to ADP-ribosylation factors. ArfGAPs are critical for cargo sorting in the Golgi-to-ER traffic. However, the role of ArfGAPs in sorting into intralumenal vesicles (ILVs) in multivesicular bodies (MVBs) in *post*-Golgi traffic remains unclear. Exosomes are extracellular vesicles (EVs) of endosomal origin. EVs mediate cell-to-cell communication, and CD63 is an EV marker. CD63 is enriched in intralumenal vesicles (ILVs) in MVBs of cells. However, the secretion of CD63 positive EVs has not been consistent with the data on CD63 localization in MVBs, and how CD63-containing EVs are formed is yet to be understood. To elucidate the mechanism of CD63 transport to ILVs, we focused on CD63 localization in MVBs and searched for the ArfGAPs involved in CD63 localization. We observed that ADAP1 and ARAP1 depletion inhibited CD63 localization to enlarged endosomes after Rab5Q79L overexpression. We tested epidermal growth factor (EGF) and CD9 localization in MVBs. We observed that ADAP1 and ARAP1 depletion affected the localization of EGF and CD9 differently. Our results indicate that there may be different populations of MVBs and that ADAP1 and ARAP1 regulate CD63 incorporation into ILVs in different MVBs.

**Summary Statement:** ADAP1 and ARAP1 regulate CD63 localization in endosomes.

## Introduction

Arf GTPase-activating proteins (ArfGAPs) mediate the hydrolysis of GTP bound to ADP-ribosylation factors (Arfs), that are small GTP-binding proteins critical for the formation of transport vesicles (Kahn et al., 2008, Donaldson and Jackson, 2011, Sztul et al., 2019). As ArfGAPs “inactivate” Arf-GTP by GTP hydrolysis, it had been thought that ArfGAPs were terminators of Arfs. However, increasing evidence indicates that ArfGAP1, the first identified and most well-studied ArfGAP, plays an important role in cargo sorting during the formation of COPI vesicles (Spang et al., 2010, Kahn, 2011, East and Kahn, 2011, Shiba and Randazzo, 2012). One binding site of COPI to Arf1-GTP binds to the cargo; therefore, the elimination of Arf1-GTP could be required before coat binding to the cargo (Arakel EC, 2019, Dodonova et al., 2017). Consistently, the roles of ArfGAPs have been studied by depleting ArfGAPs rather than overexpressing them to promote GTP hydrolysis on Arf (Shiba et al., 2013, Moshiri et al., 2017, Segeletz et al., 2018, Schoppe et al., 2020, Watanabe et al., 2021).

Extracellular vesicles (EVs) are secreted vesicles that play a role in cell-to-cell communication by transferring their cargos to other cells (Raposo and Stoorvogel, 2013, Mathieu et al., 2019, LeBleu and Kalluri, 2020). Exosomes are EVs of endosomal origin. Early endosomes are formed by the fusion of endocytic and biosynthetic vesicles and mature into multivesicular bodies (MVBs) by inward budding of the limiting membrane to form intralumenal vesicles (ILVs). MVBs fuse with the plasma membrane (PM), and ILVs are secreted as exosomes. CD63 is a tetraspanin protein known to be a marker of exosomes and is enriched in MVBs (Pols and Klumperman, 2009, Stuffers et al., 2009). CD63 forms a complex with other cargo of exosomes, such as Syntenin-1 or LMP1 (Latysheva et al., 2006, Verweij et al., 2011), therefore it could function as a cargo receptor for exosomes.

The epidermal growth factor (EGF) and its receptor (EGFR) are well-known cargo molecules transported to ILVs (Lill and Sever, 2012, Tomas et al., 2014). When EGF binds to EGFR in the plasma membrane, the EGF/EGFR complex is internalized and transported to the early endosomes. The limiting membrane of early endosomes is inwardly deformed, forming ILVs. ILV formation is mediated by the endosomal sorting complex required for transport (ESCRT) complexes, and the cytoplasmic tail of EGFR, along with the activated signaling complex, is transported into ILVs. MVBs were fused to the lysosomes, and EGF/EGFR, with the signaling complex bound to the cytoplasmic tail of EGFR, was degraded in the lysosomes.

CD63 transport to ILVs may differ from that of the EGF pathway. Quadruple depletion of ESCRT proteins changed the morphology of MVBs, but still formed MVBs where CD63 was localized in ILVs, although EGFR was observed in the limiting membrane (Stuffers et al., 2009). In addition, whether the secretion of CD63 exosome is dependent on ESCRTs is unclear. The depletion of ESCRT proteins often leads to different results: whether the secretion of CD63-positive vesicles is decreased, increased, or remains constant is dependent on different cell lines, protocols, and laboratories (Baietti et al., 2012, Matsui et al., 2021, Colombo et al., 2013).

CD9 is an exosomal marker that binds and colocalizes with EGFR (Murayama et al., 2008). CD9 localizes to the PM more often than CD63 (Mathieu et al., 2021). When CD63 is mutated to localize to the PM, the cells secrete more exosomes; therefore, secretion assays could detect vesicles of PM origin that have a similar size to those of endosomal origin (Fordjour et al., 2022). Furthermore, inhibition of CD63 localization in endosomes promotes the release of exosomes (Larios et al., 2020). The exosome fraction of the secretion assay may include EVs derived from the PM, which are similar in size to EVs of endosomal origin, although not all EVs in the exosome fraction are from the PM. Depletion of Rab27, which is involved in the secretion of secretory granules, decreases the secretion of CD63-positive exosomes (Ostrowski et al., 2010), suggesting that certain exosome fractions in the secretion assay are of endosomal origin. However, how CD63-positive vesicles are formed and whether the mechanism differs from that of CD9 or EGFR are not well understood.

To dissect the CD63 pathway, we focused on CD63 localization. We identified the ArfGAPs that regulate CD63 localization in ILVs using siRNAs targeting ArfGAPs. We used Rab5Q79L to enlarge endosomes. Rab5 is a small G protein that mediates fusion to early endosomes, and its constitutively GTP-bound form mutant, Rab5Q79L, stimulates endosomal fusion, resulting in enlarged endosomes with many ILVs (Stenmark et al., 1994, Wegner et al., 2010). Using ArfGAP siRNAs, we observed that depletion of ADAP1 and ARAP1 inhibited CD63 localization in Rab5-endosomes. We analyzed EGF and CD9 localization in ADAP1 and ARAP1KD cells. Our results indicate that ADAP1 and ARAP1 regulate CD63 localization in ILVs and differentially regulate EGF and CD9 localization.

## Results

### ADAP1 and ARAP1 were determined by siRNA-screening of ArfGAPs in MCF7 cells

To identify the ArfGAPs that regulate CD63 transport to ILVs, we used the human mammary carcinoma cell line MCF7, as we could see clear localization of CD63 inside enlarged endosomes by overexpression of Rab5Q79L. In humans, 31 ArfGAPs are encoded. Of these, AGAP4 siRNA can target AGAP4 to AGAP10 mRNAs; therefore, we used 25 siRNAs against ArfGAPs (Kahn et al., 2008, Shiba et al., 2013). We searched for siRNAs targeting ArfGAPs that decreased CD63 localization in ILVs (see Materials and Methods). We observed in ADAP1 and ARAP1 knockdown (KD) cells that CD63 localization in ILVs was decreased (Fig. 1). To confirm ADAP1 and ARAP1 depletion, we performed western blotting and observed that ADAP1 and ARAP1 were depleted by 80.4% and 96.3%, respectively, in siRNA-transfected cells (Fig. 1A and B). To quantify CD63 localization, we stained CD63 with an antibody and analyzed CD63 localization in endosomes marked with GFP-Rab5Q79L (Fig. 1C). In control cells, CD63 was localized inside the endosomes, whereas in ADAP1 and ARAP1KD cells, CD63 localization in the endosomes decreased (Fig. 1C). We quantified the number of Rab5Q79L-positive endosomes with or without CD63 and calculated the percentage of Rab5-endosomes without CD63 per cell. We observed that Rab5-endosomes without CD63 increased 20-25% in ADAP1 and ARAP1KD cells (Fig. 1D, control: 50.6%, ADAP1KD: 72.4%, ARAP1KD: 74.3%).

**Figure 1.**
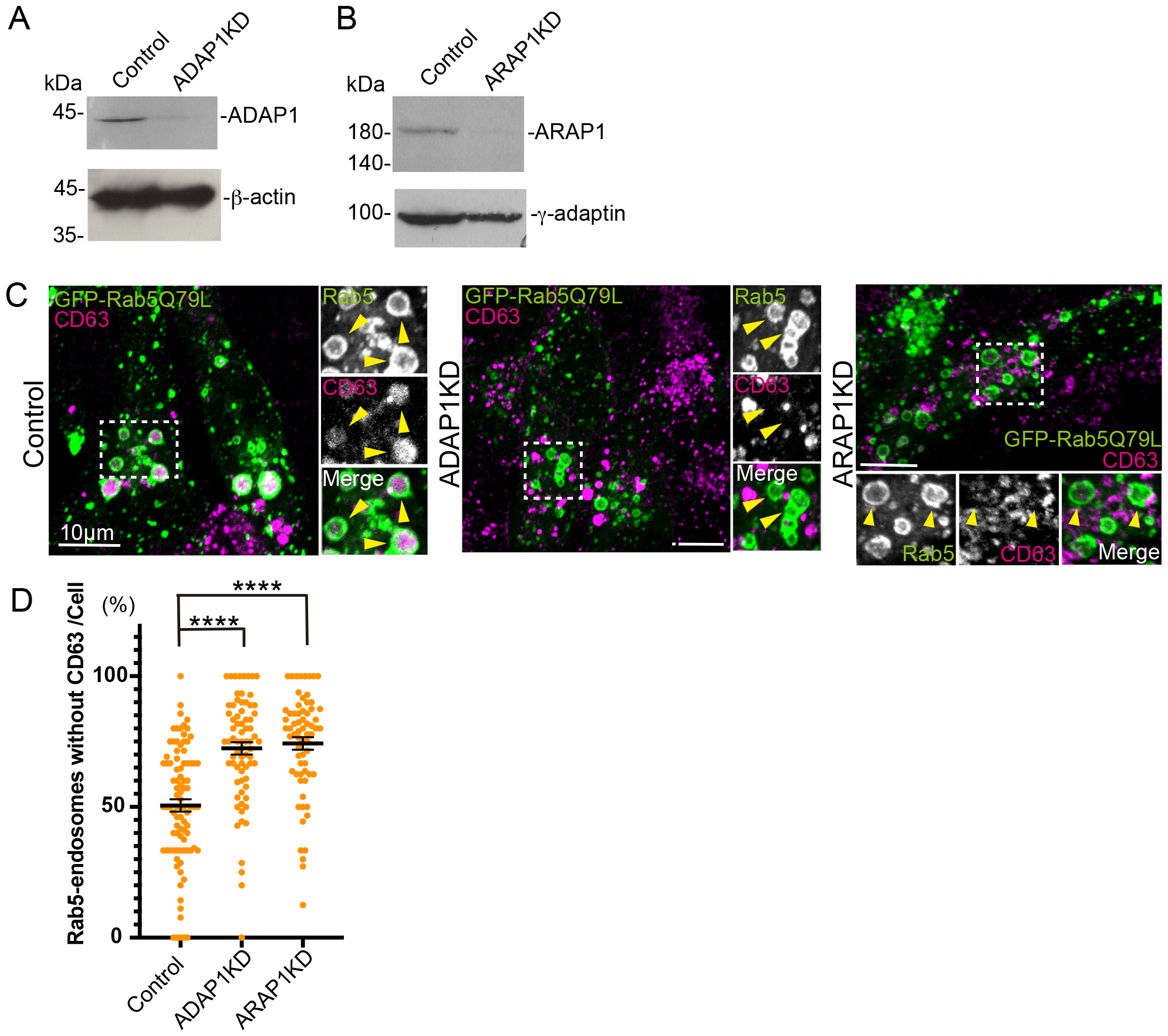
In ADAP1 and ARAP1KD cells, CD63 incorporation in endosomes is inhibited. MCF7 cells were transfected with control and ADAP1 siRNAs, and the cell lysates were subjected to western blotting. Note that ADAP1 was efficiently downregulated in ADAP1 siRNA-transfected cells compared with control siRNA-transfected cells. **B**. A similar experiment for ARAP1. Note that ARAP1 was efficiently downregulated by its siRNA. **C**. The siRNA-transfected cells were transfected with GFP-Rab5Q79L (green) and stained with anti-CD63 (red). The insets were enlarged and Rab5 positive endosomes were indicated by arrowheads. Note that CD63 was filled in endosomes in control cells, whereas CD63 signal was lost in ADAP1 and ARAP1KD cells. **D**. The Rab5-positive endosomes were classified with or without CD63, and the percentage of Rab5-endosomes without CD63 per cell was shown. Each dot represents each cell. Error bar; standard error of means (SEM). Control; *n =* 91, ADAP1KD; *n* = 73, ARAP1KD; *n =* 67. Significance was calculated by the Mann-Whitney test. ****; *p <* 0.0001.

Next, we examined whether the decrease in CD63 localization in endosomes was due to off-target effects. We transfected wild-type ADAP1 into ADAP1KD cells to determine whether the phenotype of the ADAP1KD cells was rescued (Fig. 2A). In control cells, CD63 was localized to endosomes, and in ADAP1KD cells, the number of endosomes without CD63 increased, as shown in Fig. 1. When we overexpressed ADAP1 in ADAP1KD cells, CD63 localization in the endosomes was reversed (ADAP1KD + ADAP1OE). Overexpression of ADAP1 in MCF7 cells did not affect CD63 localization in endosomes (ADAP1OE). We quantified endosomes without CD63 and observed that endosomes without CD63 were decreased by overexpression of ADAP1 in ADAP1KD cells compared to ADAP1KD cells (Fig. 2B, Control; 36.9%, ADAP1KD; 61.3%, ADAP1KD + ADAP1OE; 35.3%, ADAP1OE; 42.9%). These results confirmed that the decrease in Rab5-endosomes without CD63 in ADAP1KD cells was not an off-target effect.

**Figure 2.**
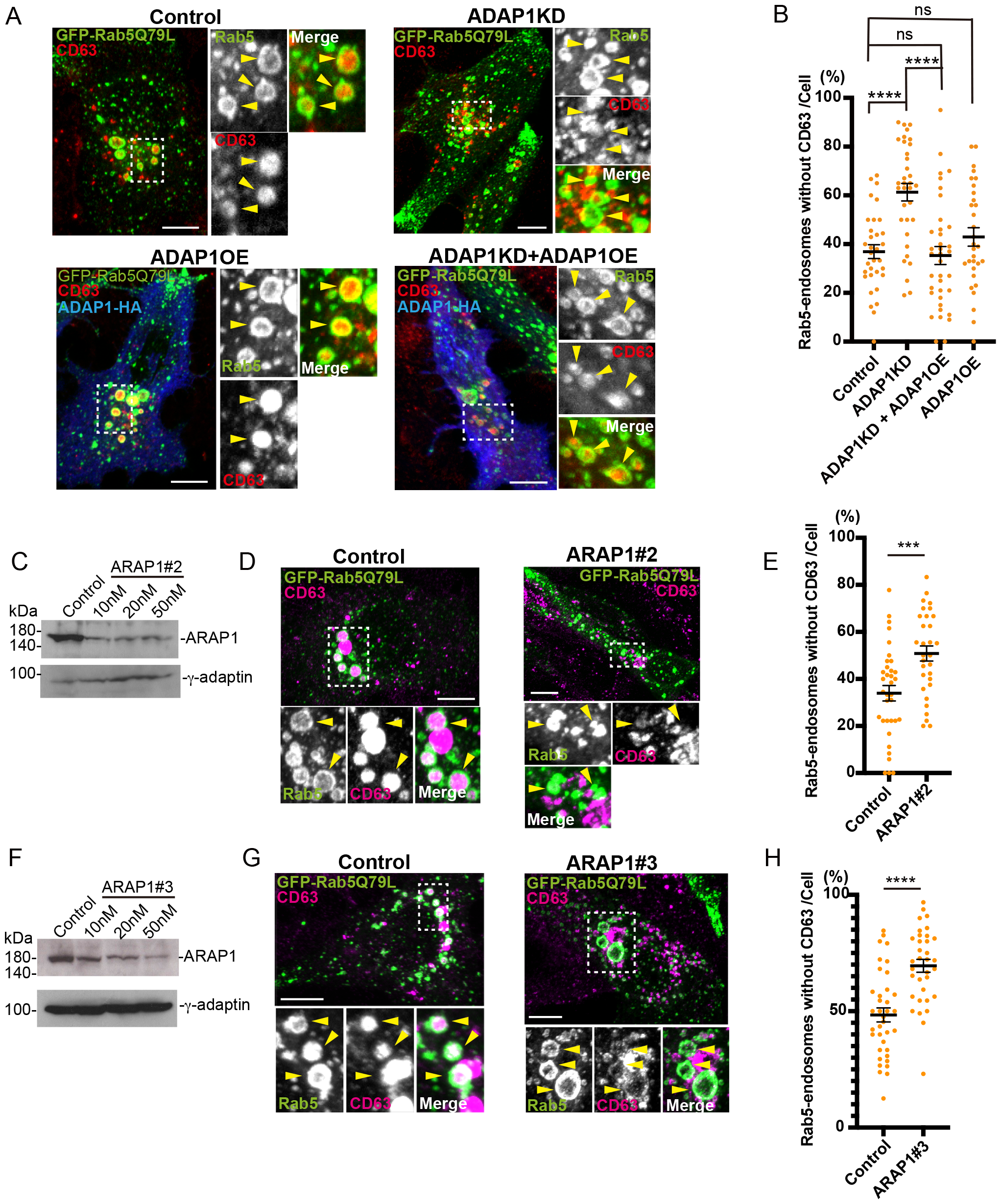
The increase in endosomes without CD63 in ADAP1 and ARAP1KD cells is not due to off-target effects. **A.** MCF7 transfected control or ADAP1 siRNA were transfected with human ADAP1-HA (ADAP1 overexpression; ADAP1OE, blue) and GFP-Rab5Q79L (green). CD63 was stained with the CD63 antibody (red). CD63 signal was seen in the control, while the signal disappeared in ADAP1KD cells. In ADAP1KD + ADAP1OE cells, the CD63 signal in the endosome was recovered. ADAP1OE cells did not show much effect on CD63 in endosomes. **B**. The results of A were quantified and the percentage of endosomes without CD63 was shown with SEM. Control; *n =* 33, ADAP1KD; *n =* 34, ADAP1KD + ADAP1OE; *n =* 37, ADAP1OE; *n =* 31. The Mann-Whitney test was performed. ****; *p <* 0.0001, *ns*; not significant. **C**. MCF7 was transfected with siRNA ARAP1#2 as indicated and subjected to western blotting. **D**. The cells were transfected with control or ARAP1#2 siRNA and immunofluorescence was performed as in A. **E**. Rab5-endosomes without CD63 was shown as in B. Control; n = 35, ARAP1#2; n = 31. The Mann-Whitney test was performed. **; *p <* 0.01. **F**. ARAP1#3 siRNA was tested for western blotting as in C. **G**. Immunofluorescence was performed for control and ARAP1#3 cells. **H**. Endosomes without CD63 were quantified. Control; n = 34, ARAP1#3; n = 37. The Mann-Whitney test was performed. ****; *p <* 0.0001.

We performed rescue experiments with ARAP1. However, overexpression of ARAP1 itself increased the number of endosomes without CD63; therefore, we could not observe a rescue of the phenotype by overexpression of ARAP1 in ARAP1KD cells (data not shown). To check the off-target effects of ARAP1 in another way, we analyzed whether two different ARAP1 siRNAs increased the number of endosomes without CD63. Our experiments typically used a mixture of four different siRNAs targeting a single gene. In this experiment, we performed western blotting on the lysates of cells transfected with 10, 20, and 50 nM of individual siRNAs, #2 and #3 targeting to ARAP1 (Fig. 2C and D). For #2 siRNA, we observed a knockdown of 81.8% at 20 nM (Fig. 2C). Therefore, we conducted immunofluorescence using 20 nM #2 siRNA and observed that CD63 localization in Rab5-endosomes was inhibited compared to that in control cells (Fig. 2D). We quantified Rab5-endosomes and observed that Rab5-endosomes without CD63 increased (Fig. 2E, control: 33.9%, ARAP1KD#2: 50.8%, *p <* 0.001). Similarly, we performed western blotting for #3 siRNA of ARAP1 and observed a 78.8% knockdown at 50 nM (Fig. 2F). Immunofluorescence analysis revealed that Rab5-endosomes without CD63 increased (Fig. 2G and H, control: 48.4%, ARAP1KD#3: 69.5%, *p <* 0.0001). These results confirmed that the perturbations in CD63 localization in Rab5-endosomes of ADAP1 and ARAP1KD cells were not due to off-target effects.

### EGF localization in ADAP1 and ARAP1 KD cells

To analyze whether EGF localization in Rab5-endosome is affected by ADAP1 and ARAP1KD cells, we internalized fluorescently labeled EGF for 5 min in HeLa cells, washed them, and incubated them for 40 min to inhibit EGF degradation under Leupeptin. In the case of EGF, its localization in endosomes was usually dotty (Fig. 3A, control). As CD63 showed 100% filling in Rab5-endosomes (Fig. 1C, control), for EGF, we classified the phenotype into 3 classes; i. endosomes ≧30% filled in; ii. < 30% endosomes filled in, which is often EGF on the limiting membranes; and iii. endosomes without EGF signals (none). For ADAP1KD cells, we observed a small increase of “endosomes ≧30% filled in”, while “endosomes < 30% filled in” and “none” were not changed (Fig. 3A and B, Control i: 39.2%, ii: 40.5%, iii: 20.2%, ADAP1KD i: 47.2%, *p* < 0.0001, ii: 37.2%, iii: 20.1%). For ARAP1KD cells, we did not observe any differences (Fig. 3C, control i: 29.7%, ii: 47.3%, iii: 23.2%: ARAP1KD i: 31.6%, ii: 45.6%, iii: 22.8%). ARAP1 was reported to be involved in EGF internalization and degradation (Yoon et al., 2008, Daniele et al., 2008, Yoon et al., 2011), however, at least for the localization of EGF in Rab5Q79L overexpressing cells, we did not see the difference in ARAP1KD cells.

**Figure 3.**
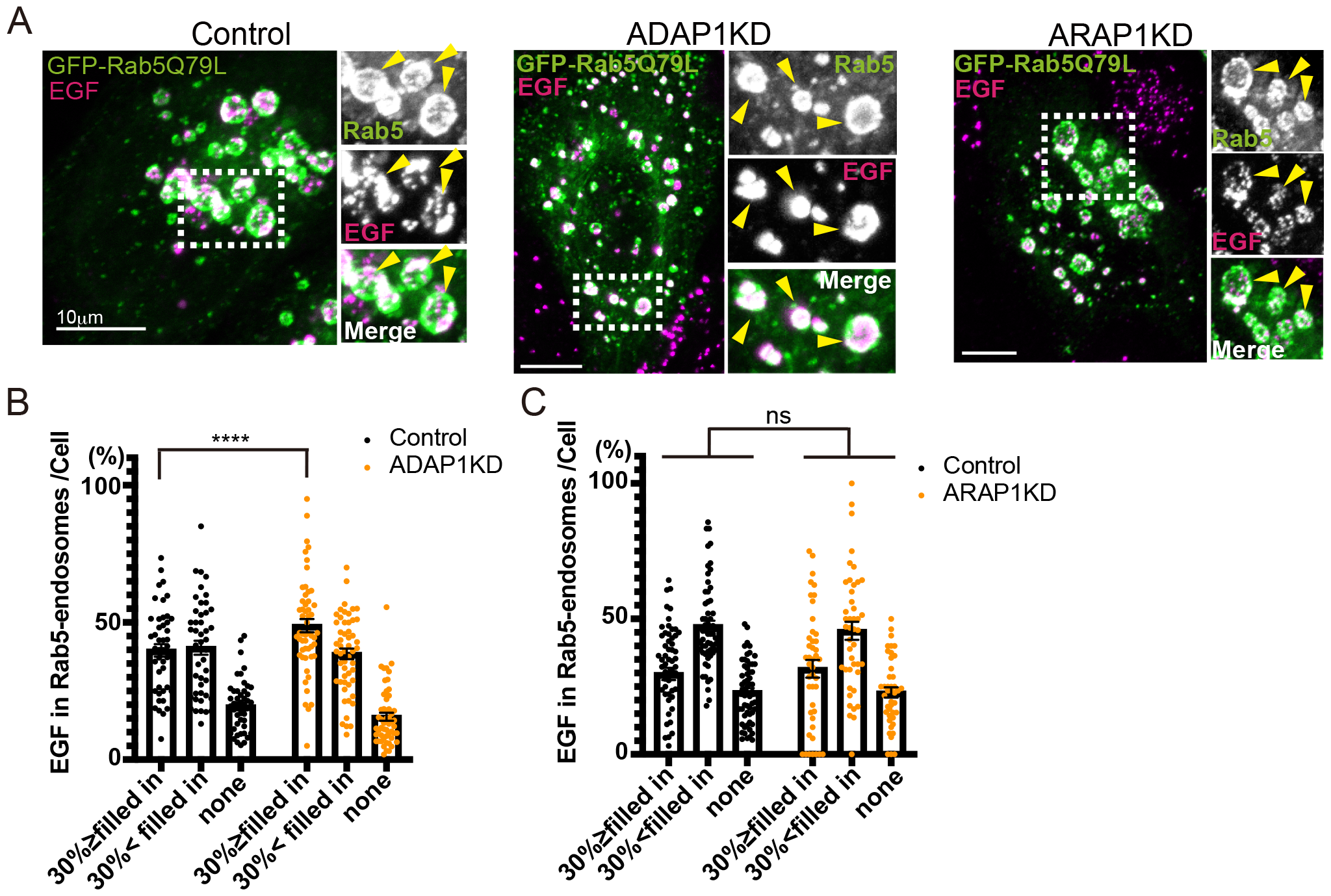
EGF localization in Rab5-endosomes is not altered in ADAP1 and ARAP1KD cells. **A.** HeLa cells were transfected with siRNAs as indicated and EGF-Alexa 555 was internalized in HeLa cells for 5 min, washed, and incubated for 40 min under leupeptin. Note that EGF was in endosomes but not 100% filled, for control, ADAP1, and ARAP1KD cells. **B**. In control and ADAP1KD cells, Rab5-endosomes with EGF were classified as indicated, and the percentages of each endosome per cell are shown with SEM. Control; *n =* 46, ADAP1KD; *n =* 53. Significance was tested using Mixed-effects analysis. *ns*; not significant. **C**. Rab5-endosomes in control and ARAP1KD cells were analyzed as in B. Control; *n =* 60, ARAP1KD; *n =* 44. Significance was tested using the Mixed-effects analysis. *ns*; not significant.

Our results indicate that in ADAP1 and ARAP1KD cells, EGF localization in Rab5-endosomes was not inhibited.

### CD9 localization in Rab5-endosomes

Next, we analyzed CD9 localization in ADAP1 and ARAP1KD cells. CD9 showed a more prominent signal in the plasma membrane than CD63, and an endosome signal (Fig. 4A and Fig. 1C). In the case of CD9, many endosomes were filled partially as well as that were filled 100%. Therefore, we classified endosomes into three classes: EGF, i: 100% filled in, ii: partially filled in, and iii: no CD9. We quantified endosomes and calculated the percentage of Rab5-endosomes in each category per cell. We observed that in ADAP1 and ARAP1KD cells, Rab5-endosomes without CD9 were increased (Fig. 4B). The endosomal phenotype was different; in ADAP1KD cells, endosomes partially filled with CD9 were decreased (Fig. 4 B, Control, i: 24.0%, ii: 34.6%, iii: 41.3%, ADAP1KD, I: 19.0%, ii: 23.8%, *p <* 0.05; iii: 53.1%, *p <* 0.01), whereas in ARAP1KD cells, endosomes 100% filled with CD9 were decreased compared with control cells (Fig. 4B, ARAP1KD, I: 10.8%, *p <* 0.05, ii: 27.2%, iii: 61.9%, *p <* 0.0001). These results suggest that ADAP1 and ARAP1 affect CD9 localization in ILVs.

**Figure 4.**
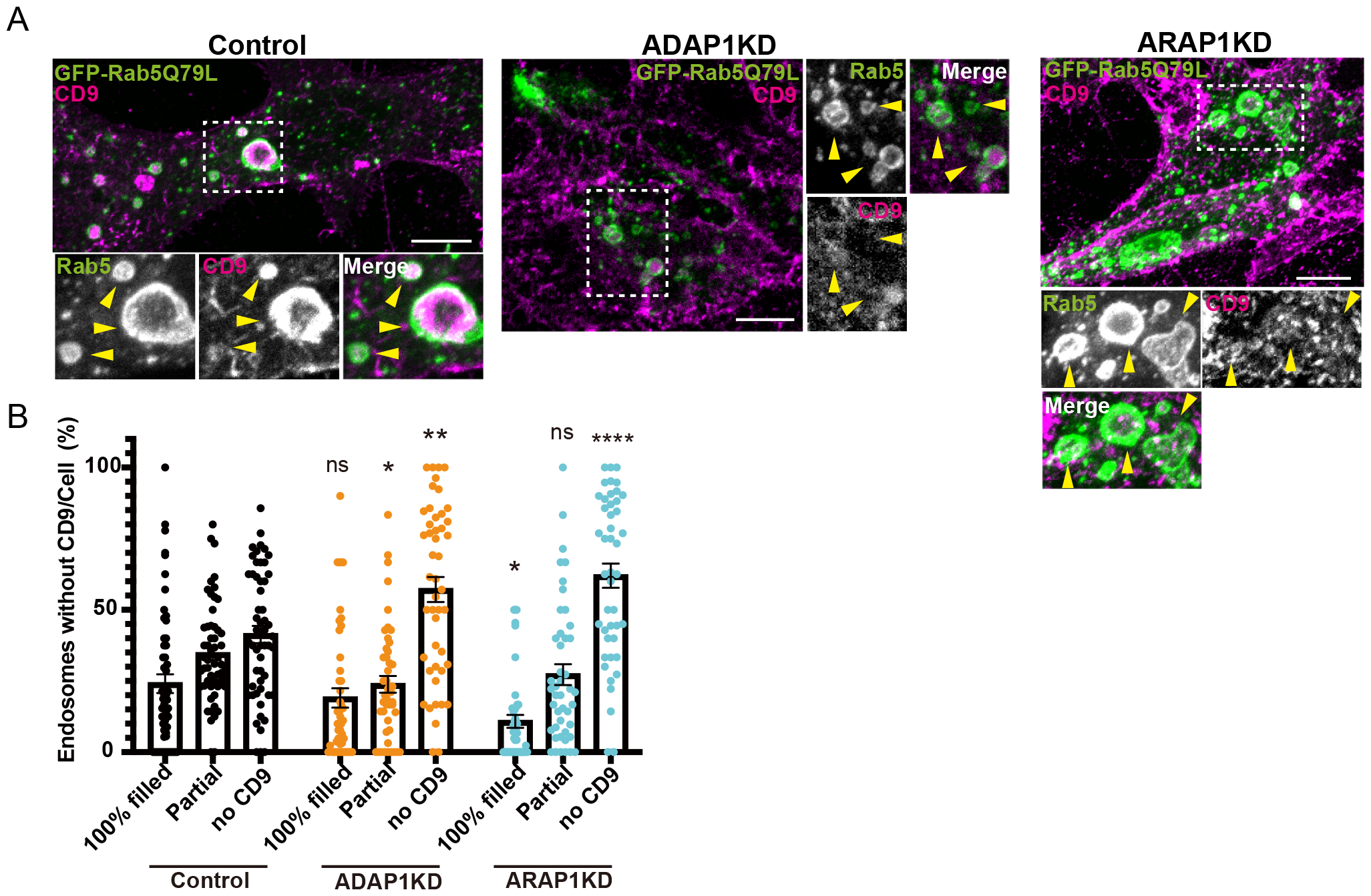
The incorporation of CD9 in Rab5-endosomes is inhibited in ADAP1 and ARAP1KD cells. **A.** MCF7 cells were transfected with each siRNA as indicated, then transfected with GFP-Rab5Q79L (green), and stained with anti-CD9 antibody (red). CD9 is localized in endosomes in control cells, while in ADAP1 and ARAP1KD cells, CD9 localization in endosomes was decreased. **B**. The endosomes were classified as indicated and the percentages of each cell were shown with SEM. Control; *n =*55, ADAP1KD; *n =* 47, ARAP1KD; *n =* 45. Significance was tested using mixed-effects analysis and Sidak’s multiple comparison test. *; *p <* 0.05, **; *p <* 0.01, ***; *p <* 0.001, ****; *p <* 0.0001, *ns*; not significant.

## Discussion

In this study, we identified ADAP1 and ARAP1 as ArfGAPs that promote CD63 localization in ILVs. We observed that ADAP1 and ARAP1 affect CD9 localization in ILVs, whereas EGF localization in ILVs was not perturbed, rather, even EGF localization in endosomes was increased in ADAP1KD cells.

EGF and CD63 transport pathways to ILVs have been reported to differ. EGF is known to be dependent on the ESCRT complex, whereas CD63 has an ESCRT-independent pathway to localize to ILVs (Stuffers et al., 2009, van Niel et al., 2011, Edgar et al., 2014). Our results align with previous studies. In ADAP1 and ARAP1KD cells, inhibition of CD63 localization in ILVs was observed (Fig. 1 and 2). However, EGF localization in ILVs was not inhibited. In ADAP1KD cells, there was a slight increase in the extent of “endosomes ≧30% filled in” (Fig. 3B). ARAP1 has been known to be involved in EGF transport. In ARAP1 KD cells, EGF internalization was inhibited by FACS analysis (Yoon et al., 2008). Two reports have suggested that EGFR degradation is accelerated in ARAP1KD cells (Yoon et al., 2008, Yoon et al., 2011), whereas another report suggested that EGFR degradation is inhibited in ARAP1KD cells (Daniele et al., 2008). Either way, ARAP1 would be involved in EGFR trafficking, but in our analysis, at least EGF localization in Rab5-endosomes was not perturbed in ARAP1KD cells.

CD9 is known to bind to EGFR and colocalize with EGFR (Murayama et al., 2008) and has been observed in CD63-positive vesicles (Matsui et al., 2021, Fordjour et al., 2022), though intracellular distribution of CD9 is different than CD63 (Maeda et al., 2023, Mathieu et al., 2021). We observed that in ADAP1 and ARAP1KD cells, the incorporation of CD9 into Rab5-endosomes was inhibited. The pattern of inhibition was different; in ADAP1KD cells, “endosomes with partially filled in” was inhibited, whereas in ARAP1KD cells, “endosomes 100% filled in” was inhibited (Fig. 4B). We speculate that there are two different MVBs (Fig. 5; MVB1 and MVB2). CD63-positive ILVs have been reported to be smaller (50 nm in diameter) than ILVs formed by EGF stimulation (60 nm) (Edgar et al., 2014). We hypothesized that MVB1 has ILVs for EGFR, and MVB2 does not (Fig. 5). The depletion of ADAP1 inhibits ILVs for CD63 and CD9 of “endosomes partially filled”. Therefore, it could stimulate more EGFR ILVs in MVB1, leading to a small increase in EGF localization in “endosomes ≧30% filled in (Fig. 3B). In ARAP1KD cells, EGF localization was not affected (Fig. 4B); therefore, MVB2 is independent of EGF, and ARAP1 regulates the ILVs of CD63 and CD9 of “endosomes 100% filled in” for MVB2. MVB1 does not seem to mature into MVB2, or vice versa. If MVB1 matures into MVB2, the depletion of ADAP1 should have decreased “endosomes with 100% filled” with CD9 as well. However, our results did not show such phenomena, rather, we observed ADAP1 depletion increased endosomes without CD9 (Fig. 4B). Therefore, ADAP1 and ARAP1 may regulate different MVBs.

**Figure 5.**
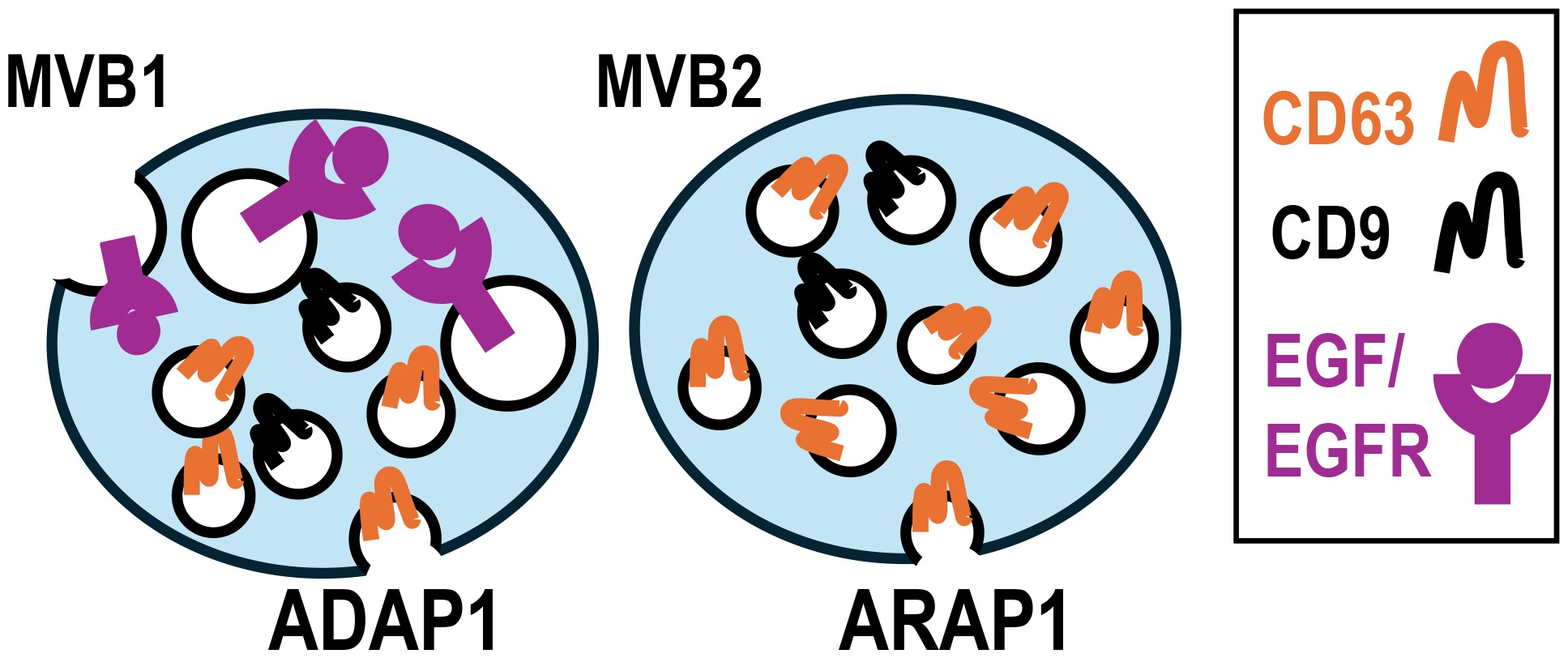
ADAP1 and ARAP1 regulate different MVBs. ADAP1 and ARAP1 function in CD63 localization in MVBs. At least two different MVBs were observed: MVB1 and MVB2. In this model, MVB1 cells contained ILVs for EGF/EGFR. ADAP1 regulates ILVs of CD63 in MVB1, and ARAP1 regulates ILVs of CD63 in MVB2. ILVs of CD9 were observed in both MVB1 and MVB2.

ARAP1 exhibits GAP activity against Arf1 and Arf5 dependent on PI (3,4,5) P3 (Miura et al., 2002). In contrast, ADAP1 exhibits GAP activity against Arf6 (Venkateswarlu et al., 2004, Duellberg et al., 2021). Future studies should investigate how ADAP1 and ARAP1 affect exosome secretion and the mechanisms by which ADAP1 and ARAP1 differentially regulate CD63 localization in MVBs. In this study, we found that ADAP1 and ARAP1 play important roles in the localization of CD63. Our work contributes to the understanding of the heterogeneity of MVBs and exosomes.

## Materials and Methods

### Reagents

ON-TARGET plus non-targeting siRNA #4, the siGENOME SMART pool of human ADAP1 and ARAP1, #2 and #3 siRNAs of ARAP1, and human ADAP1 and ARAP1 plasmids were purchased from Horizon Discovery Ltd. (Cambridge, UK). N-terminal FLAG-tagged human ARAP1 (Miura et al., 2002) and canine GFP-Rab5Q79L were kindly provided by Dr. Paul Randazzo (NIH, USA) and Dr. Marino Zerial (Max Planck Institute of Molecular Cell Biology and Genetics, Germany), respectively. ADAP1 was inserted into pcDNA3.1, with the C-terminally tagged using EcoRI and XhoI. The mouse monoclonal anti-human CD63 antibody (H5C6, DSHB, IA) was kindly provided by Dr. Eiji Morita (Hirosaki University, Japan). The rabbit polyclonal anti-ARAP1 was procured from Abcam (Cambridge, UK), rat monoclonal anti-HA (3F10) from Merck (Darmstadt, Germany), mouse monoclonal anti-myc (9E10) from Invitrogen (Waltham, MA, USA), rabbit polyclonal anti-FLAG from Merck (Darmstadt, Germany), mouse monoclonal anti-β-actin (8H10D10) from Cell Signaling (Danvers, MA, USA), and mouse monoclonal anti-Adaptin γ (88) from BD Biosciences (Franklin Lakes, NJ, USA). Secondary antibodies of goat anti-Rabbit IgG conjugated with Alexa Fluor 568 and anti-Mouse Alexa conjugated with Fluor 594 were purchased from Thermo Fisher Scientific (Waltham, MA, USA), donkey Anti-Rabbit conjugated with Alexa Fluor^®^ 488 from Abcam, and AffinPure donkey anti-mouse and rabbit IgG conjugated with peroxidase from Jackson ImmunoResearch (West Grove, PA, USA).

### Cell culture

The human mammary carcinoma cell line, MCF7, was purchased from RIKEN BRC (Tsukuba, Japan). MCF7 cells were maintained in DMEM (Nacalai Tesque, Kyoto, Japan) supplemented with 10% FBS (Regular; CORNING, NY, USA), 1x MEM Non-essential Amino Acids Solution (Wako, Osaka, Japan), and 1 mmol/L Sodium Pyruvate Solution (Wako). Human cervical cancer HeLa cells were purchased from RIKEN BRC. HeLa cells were maintained in DMEM supplemented with 10% FBS. Cells were typically used by passage six.

### siRNA screening of ArfGAPs in MCF7 cells

We tested several cells that clearly showed CD63 localization in Rab5-endosomes and decided to use MCF7 overexpressing myc-tagged CD63 cells (kindly provided by Dr. Eiji Morita). A custom cherry-pick siRNA library was constructed from 25 human ArfGAPs and control siRNAs using Horizon Discovery Ltd. (Cambridge, UK). Knockdown efficiency was confirmed by ArfGAP3 siRNA using western blotting, and almost 100% knockdown was achieved. After 24 h of siRNA transfection, the medium was changed, and the cells were transfected with GFP-Rab5Q79L and CD63-myc using Lipofectamine 3000, according to the manufacturer’s protocol (Thermo Fisher Scientific, MA, USA). The medium was changed after 24 h and the cells were incubated for another 24 h. The siRNA transfection duration was 72 h. The cells were fixed with 4% PFA/PBS and processed for immunofluorescence analysis. A confocal microscope (Nikon C2, Tokyo, Japan) was used to capture images using 60× objective (NA1.40) and 100× objective (NA 1.45). We captured images of Rab5Q79L-overexpressing cells randomly for each siRNA using the same settings of confocal microscope. The experiments were repeated twice and more than 30 cells were analyzed, except for ACAP3. Thirteen ArfGAPs were found to increase the endosomes without CD63 expression. We repeated the screening, including the siRNA of SMCR8, which was recently reported to be an ArfGAP (Su et al., 2021) with CD63 antibody for endogenous CD63. Cells transfected with the four ArfGAP siRNAs still showed an increase in endosomes without CD63. Among the four ArfGAPs, we confirmed ADAP1 and ARAP1 for knockdown of their proteins and a decrease in CD63 localization in endosomes (Fig. 1).

### Immunofluorescence

A total of 5 × 10^4^ MCF7 cells were seeded on coverslips and transfected with siRNAs (10 nM) using Lipofectamine RNAiMAX (Thermo Fisher Scientific, MA, USA) by reverse transfection according to the manufacturer’s protocol. The medium was replaced or cells were transfected with plasmids of GFP-Rab5Q79L 24 h after transfection using Lipofectamine LTX (Thermo Fisher Scientific, MA, USA) according to the manufacturer’s protocol. Four hours after the plasmid transfection, the medium was changed. Cells were incubated for 72 h after siRNA transfection.

HeLa cells were used for EGF internalization. After 68 h of siRNA transfection, cells were treated with 100 mg/ml leupeptin (Peptide Institute, Inc. Osaka, Japan). During leupeptin treatment, cells were serum-starved for 1 h, incubated with 100 ng/ml EGF-Alexa 555 (Thermo Fisher Scientific, MA, USA) for 5 min, washed, and incubated at 37 °C for 40 min. Treatment with leupeptin was performed for 4 h.

The cells were fixed with 4% paraformaldehyde (PFA) at room temperature for 15 min. After permeabilization and blocking with 0.02% saponin and 0.2% BSA/PBS at room temperature for 30 min, cells were stained with primary antibodies in 0.02% saponin and BSA/PBS for 2 h. After washing three times in 0.02% saponin/0.2% BSA/PBS, the cells were stained with secondary antibodies in PBS for 45 min and washed thrice with PBS. Cells were mounted using Mowiol (Calbiochem, Darmstadt, Germany). We used confocal microscope (Nikon C2 and AX/AXR, Tokyo, Japan) and captured images using 60× objective (NA1.40 and NA1.42) and 100× objective (NA 1.45).

### Image analysis and statistics

For quantification, the same confocal microscopy settings were used for control and samples. We randomly captured an area of GFP-Rab5Q79L expressing cells and counted the number of endosomes with and without CD63. The percentage of endosomes without CD63 relative to the total number of endosomes per cell was calculated. The experiments were repeated at least 3 times and at least 30 cells were counted. Statistical analyses were performed using GraphPad PRISM version 10 (GraphPad Software, San Diego, California, USA; www.graphpad.com).

### Western blotting

A total of 5 × 10^4^ MCF7 cells were transfected with siRNAs using Lipofectamine RNAiMAX. After 72 hr of transfection, the cells were washed twice with PBS and lysed with 50 μL of sample buffer (50 mM Tris-HCl pH 6.4, 8% Glycerol, 0.1% bromophenol blue, 50 mM DTT, 2% SDS). An aliquot of 30 μL from each sample was subjected to electrophoresis on 8% or 15% polyacrylamide gel for ARAP1 and ADAP1 each and transferred into an Immobilon^®^-P transfer membrane (Merck Millipore Ltd., Germany). Membranes were blocked with 5% skim milk (Nacalai Tesque, Japan) in Tris-buffered saline containing 0.1% Tween 20 (TBS-T). The membranes were incubated with primary antibodies diluted in Signal Enhancer HIKARI Solution A (Nacalai Tesque, Japan), washed with TBS-T 3 times, and incubated with horseradish peroxidase-conjugated secondary antibodies for 45 min. Proteins were visualized by enhanced chemiluminescence using an ECL™ western blotting Analysis System (Cytiva, Marlborough, MA, USA). Quantification was performed using the imageJ Fiji software (Schindelin et al., 2012).

## Acknowledgments

We greatly thank Dr. Eiji Morita for providing reagents and helpful discussions.

## Competing interests

No competing interests declared.

## Funding

This project was supported by an Iwate University grant to YS.

## Notes

### Competing Interest Statement

The authors have declared no competing interest.

## References

Arakel, E. C., Huranova, M., Estrada, A. F., Rau, E. M., Spang, A., and Schwappach, B. (2019). Dissection of GTPase-activating proteins reveals functional asymmetry in the COPI coat of budding yeast. Journal of cell science, 132.

Baietti, M. F., Zhang, Z., Mortier, E., Melchior, A., Degeest, G., Geeraerts, A., Ivarsson, Y., Depoortere, F., Coomans, C., Vermeiren, E., Zimmermann, P., and David, G. (2012). Syndecan-syntenin-ALIX regulates the biogenesis of exosomes. Nature cell biology, 14, 677–685.

Colombo, M., Moita, C., van Niel, G., Kowal, J., Vigneron, J., Benaroch, P., Manel, N., Moita, L. F., Théry, C., and Raposo, G. (2013). Analysis of ESCRT functions in exosome biogenesis, composition and secretion highlights the heterogeneity of extracellular vesicles. Journal of cell science, 126, 5553–5565.

Daniele, T., Di Tullio, G., Santoro, M., Turacchio, G., and De Matteis, M. A. (2008). ARAP1 regulates EGF receptor trafficking and signalling. Traffic, 9, 2221–2235.

Dodonova, S. O., Aderhold, P., Kopp, J., Ganeva, I., Röhling, S., Hagen, W. J. H., Sinning, I., Wieland, F., and Briggs, J. A. G. (2017). 9Å structure of the COPI coat reveals that the Arf1 GTPase occupies two contrasting molecular environments. eLife, 6, e26691.

Donaldson, J. G., and Jackson, C. L. (2011). ARF family G proteins and their regulators: roles in membrane transport, development and disease. Nature reviews. Molecular cell biology, 12, 362–375.

Duellberg, C., Auer, A., Canigova, N., Loibl, K., & Loose, M. (2021). In vitro reconstitution reveals phosphoinositides as cargo-release factors and activators of the ARF6 GAP ADAP1. Proc Natl Acad Sci USA, 118.

East, M. P. and Kahn, R. A. (2011). Models for the functions of Arf GAPs. Semin Cell Dev Biol, 22, 3–9.

Edgar, J. R., Eden, E. R. and Futter, C. E. (2014). Hrs- and CD63-dependent competing mechanisms make different sized endosomal intraluminal vesicles. Traffic, 15, 197–211.

Fordjour, F. K., Guo, C., Ai, Y., Daaboul, G. G. and Gould, S. J. (2022). A shared, stochastic pathway mediates exosome protein budding along plasma and endosome membranes. J Biol Chem, 298, 102394.

Kahn, R. A. (2011). GAPs: Terminator versus effector functions and the role(s) of ArfGAP1 in vesicle biogenesis. Cell Logist, 1, 49–51.

Kahn, R. A., Bruford, E., Inoue, H., Logsdon, J. M., JR., Nie, Z., Premont, R. T., Randazzo, P. A., Satake, M., Theibert, A. B., Zapp, M. L. and Cassel, D. (2008). Consensus nomenclature for the human ArfGAP domain-containing proteins. J Cell Biol, 182, 1039–1044.

Larios, J., Mercier, V., Roux, A. and Gruenberg, J. (2020). ALIX- and ESCRT-III-dependent sorting of tetraspanins to exosomes. J Cell Biol, 219.

Latysheva, N., Muratov, G., Rajesh, S., Padgett, M., Hotchin, N. A., Overduin, M. and Berditchevski, F. (2006). Syntenin-1 is a new component of tetraspanin-enriched microdomains: mechanisms and consequences of the interaction of syntenin-1 with CD63. Mol Cell Biol, 26, 7707–18.

Lebleu, V. S. and Kalluri, R. (2020). Exosomes as a Multicomponent Biomarker Platform in Cancer. Trends Cancer, 6, 767–774.

Lill, N. L. and Sever, N. I. (2012). Where EGF receptors transmit their signals. Sci Signal, 5, pe41.

Maeda, K., Goto, S., Miura, K., Saito, K. and Morita, E. (2023). The Incorporation of Extracellular Vesicle Markers Varies Among Vesicles with Distinct Surface Charges. J Biochem.

Mathieu, M., Martin-Jaular, L., Lavieu, G. and Thery, C. (2019). Specificities of secretion and uptake of exosomes and other extracellular vesicles for cell-to-cell communication. Nat Cell Biol, 21, 9–17.

Mathieu, M., Nevo, N., Jouve, M., Valenzuela, J. I., Maurin, M., Verweij, F. J., Palmulli, R., Lankar, D., Dingli, F., Loew, D., Rubinstein, E., Boncompain, G., Perez, F. and Thery, C. (2021). Specificities of exosome versus small ectosome secretion revealed by live intracellular tracking of CD63 and CD9. Nat Commun, 12, 4389.

Matsui, T., Osaki, F., Hiragi, S., Sakamaki, Y. & Fukuda, M. (2021). ALIX and ceramide differentially control polarized small extracellular vesicle release from epithelial cells. EMBO Rep, 22, e51475.

Miura, K., Jacques, K. M., Stauffer, S., Kubosaki, A., Zhu, K., HIirsch, D. S., Resau, J., Zheng, Y. and Randazzo, P. A. (2002). ARAP1: a point of convergence for Arf and Rho signaling. Mol Cell, 9, 109–19.

Moshira, A., Humpal, D., Leonard, B. C., Imai, D. M., Tham, A., Bower, L., Clary, D., Glaser, T. M., Lloyd, K. C. and Murphy, C. J. (2017). Arap1 Deficiency Causes Photoreceptor Degeneration in Mice. Invest Ophthalmol Vis Sci, 58, 1709–1718.

Murayama, Y., Shinomura, Y., Oritani, K., Miyagawa, J., Yoshida, H., Nishida, M., Katsube, F., Shiraga, M., Miyasaki, T., Nakamoto, T., Tsutsui, S., Tamura, S., Higashiyama, S., Shimomura, I. and Hayashi, N. (2008). The tetraspanin CD9 modulates epidermal growth factor receptor signaling in cancer cells. J Cell Physiol, 216, 135–43.

Ostrowski, M., Carmo, N. B., Krumeich, S., Fanget, I., Raposo, G., Savina, A., Moita, C. F., Schauer, K., Hume, A. N., Freitas, R. P., Goud, B., Benaroch, P., Hacohen, N., Fukuda, M., Desnos, C., Seabra, M. C., Darchen, F., Amigorena, S., Moita, L. F. and Thery, C. (2010). Rab27a and Rab27b control different steps of the exosome secretion pathway. Nat Cell Biol, 12, 19–30; sup pp 1–13.

Pols, M. S. and Klumperman, J. (2009). Trafficking and function of the tetraspanin CD63. Exp Cell Res, 315, 1584–92.

Raposo, G. and Stoorvogel, W. (2013). Extracellular vesicles: exosomes, microvesicles, and friends. J Cell Biol, 200, 373–83.

Schoppe, J., Mari, M., Yavavli, E., Auffarth, K., Cabrera, M., Walter, S., Frohlich, F. and Ungermann, C. (2020). AP-3 vesicle uncoating occurs after HOPS-dependent vacuole tethering. EMBO J, 39, e105117.

Segeletz, S., Danglot, L., Galli, T. and Hoflack, B. (2018). ARAP1 Bridges Actin Dynamics and AP-3-Dependent Membrane Traffic in Bone-Digesting Osteoclasts. iScience, 6, 199–211.

Shiba, Y., Kametaka, S., Waguri, S., Presley, J. F. and Randazzo, P. A. (2013). ArfGAP3 regulates the transport of cation-independent mannose 6-phosphate receptor in the post-Golgi compartment. Curr Biol, 23, 1945–51.

Shiba, Y. and Randazzo, P. A. (2012). ArfGAP1 function in COPI mediated membrane traffic: currently debated models and comparison to other coat-binding ArfGAPs. Histol Histopathol, 27, 1143–1153.

Spang, A., Shiba, Y. and Randazzo, P. A. (2010). Arf GAPs: gatekeepers of vesicle generation. FEBS Lett, 584, 2646–2651.

Stenmark, H., Parton, R. G., Steele-Mortimer, O., Lutcke, A., Gruenberg, J. and Zerial, M. (1994). Inhibition of rab5 GTPase activity stimulates membrane fusion in endocytosis. EMBO J, 13, 1287–96.

Stuffers, S., Sem Wegner, C., Stenmark, H. and Brech, A. (2009). Multivesicular endosome biogenesis in the absence of ESCRTs. Traffic, 10, 925–37.

Su, M. Y., Fromm, S. A., Remis, J., Toso, D. B. and Hurley, J. H. (2021). Structural basis for the ARF GAP activity and specificity of the C9orf72 complex. Nat Commun, 12, 3786.

Sztul, E., Chen, P. W., Casanova, J. E., Cherfils, J., Dacks, J. B., Lambright, D. G., Lee, F. S., Randazzo, P. A., Santy, L. C., Schurmann, A., Wilhelmi, I., Yohe, M. E. and Kahn, R. A. (2019). ARF GTPases and their GEFs and GAPs: concepts and challenges. Mol Biol Cell, 30, 1249–1271.

Tomas, A., Futter, C. E. and Eden, E. R. (2014). EGF receptor trafficking: consequences for signaling and cancer. Trends Cell Biol, 24, 26–34.

Van Niel, G., Charrin, S., Simoes, S., Romao, M., Rochin, L., Saftig, P., Marks, M. S., Rubinstein, E. and Raposo, G. (2011). The tetraspanin CD63 regulates ESCRT-independent and -dependent endosomal sorting during melanogenesis. Dev Cell, 21, 708–21.

Venkateswarlu, K., Brandom, K. G. and Lawrence, J. L. (2004). Centaurin-alpha1 is an in vivo phosphatidylinositol 3,4,5-trisphosphate-dependent GTPase-activating protein for ARF6 that is involved in actin cytoskeleton organization. J Biol Chem, 279, 6205–8.

Verweij, F. J., Van Eijndhoven, M. A., Hopmans, E. S., Vendrig, T., Wurdinger, T., Cahir-Mcfarland, E., Kkieff, E., Geerts, D., Van der Kant, R., Neefjes, J., Middeldorp, J. M. and Pegtel, D. M. (2011). LMP1 association with CD63 in endosomes and secretion via exosomes limits constitutive NF-kappaB activation. EMBO J, 30, 2115–29.

Watanabe, A., Hataida, H., Inoue, N., Kamon, K., Bbaba, K., Sasaki, K., Kimura, R., Sasaki, H., Eura, Y., Ni, W. F., Shibasaki, Y., Waguri, S., Kokame, K. and Shiba, Y. (2021). Arf GTPase-activating proteins SMAP1 and AGFG2 regulate the size of Weibel-Palade bodies and exocytosis of von Willebrand factor. Biol Open, 10.

Wegner, C. S., Malerod, L., Pedersen, N. M., Prodiga, C., Bakke, O., Stenmark, H. and Brech, A. (2010). Ultrastructural characterization of giant endosomes induced by GTPase-deficient Rab5. Histochem Cell Biol, 133, 41–55.

Yoon, H. Y., Kales, S. C., Luo, R., Lipkowitz, S. and Randazzo, P. A. (2011). ARAP1 association with CIN85 affects epidermal growth factor receptor endocytic trafficking. Biol Cell, 103, 171–84.

Yoon, H. Y., Lee, J. S. and Randazzo, P. A. (2008). ARAP1 regulates endocytosis of EGFR. Traffic, 9, 2236–52.

Arakel EC, M. H., Alejandro F. Estrada, E-Ming Rau, Anne Spang, and Blanche Schwappach 2019. Dissection of GTPase-activating proteins reveals functional asymmetry in the COPI coat of budding yeast. Journal of Cell Science, 132.

Baietti, M. F., Zhang, Z., Mortier, E., Melchior, A., Degeest, G., Geeraerts, A., Ivarsson, Y., Depoortere, F., Coomans, C., Vermeiren, E., Zimmermann, P. & David, G. 2012. Syndecan-syntenin-ALIX regulates the biogenesis of exosomes. Nat Cell Biol, 14, 677–85.

Colombo, M., Moita, C., Van Niel, G., Kowal, J., Vigneron, J., Benaroch, P., Manel, N., Moita, L. F., Thery, C. & Raposo, G. 2013. Analysis of ESCRT functions in exosome biogenesis, composition and secretion highlights the heterogeneity of extracellular vesicles. J Cell Sci, 126, 5553–65.

Daniele, T., Di Tullio, G., Santoro, M., Turacchio, G. & De Matteis, M. A. 2008. ARAP1 regulates EGF receptor trafficking and signalling. Traffic, 9, 2221–35.

Dodonova, S. O., Aderhold, P., Kopp, J., Ganeva, I., Rohling, S., Hagen, W. J. H., Sinning, I., Wieland, F. & Briggs, J. A. G. 2017. 9A structure of the COPI coat reveals that the Arf1 GTPase occupies two contrasting molecular environments. Elife, 6.

Donaldson, J. G. & Jackson, C. L. 2011. ARF family G proteins and their regulators: roles in membrane transport, development and disease. Nat Rev Mol Cell Biol, 12, 362–375.

Duellberg, C., Auer, A., Canigova, N., Loibl, K. & Loose, M. 2021. In vitro reconstitution reveals phosphoinositides as cargo-release factors and activators of the ARF6 GAP ADAP1. Proc Natl Acad Sci U S A, 118.

East, M. P. & Kahn, R. A. 2011. Models for the functions of Arf GAPs. Semin Cell Dev Biol, 22, 3–9.

Edgar, J. R., Eden, E. R. & Futter, C. E. 2014. Hrs- and CD63-dependent competing mechanisms make different sized endosomal intraluminal vesicles. Traffic, 15, 197–211.

Fordjour, F. K., Guo, C., Ai, Y., Daaboul, G. G. & Gould, S. J. 2022. A shared, stochastic pathway mediates exosome protein budding along plasma and endosome membranes. J Biol Chem, 298, 102394.

Kahn, R. A. 2011. GAPs: Terminator versus effector functions and the role(s) of ArfGAP1 in vesicle biogenesis. Cell Logist, 1, 49–51.

Kahn, R. A., Bruford, E., Inoue, H., Logsdon, J. M., Jr., Nie, Z., Premont, R. T., Randazzo, P. A., Satake, M., Theibert, A. B., Zapp, M. L. & Cassel, D. 2008. Consensus nomenclature for the human ArfGAP domain-containing proteins. J Cell Biol, 182, 1039–1044.

Larios, J., Mercier, V., Roux, A. & Gruenberg, J. 2020. ALIX- and ESCRT-III-dependent sorting of tetraspanins to exosomes. J Cell Biol, 219.

Latysheva, N., Muratov, G., Rajesh, S., Padgett, M., Hotchin, N. A., Overduin, M. & Berditchevski, F. 2006. Syntenin-1 is a new component of tetraspanin-enriched microdomains: mechanisms and consequences of the interaction of syntenin-1 with CD63. Mol Cell Biol, 26, 7707–18.

Lebleu, V. S. & Kalluri, R. 2020. Exosomes as a Multicomponent Biomarker Platform in Cancer. Trends Cancer, 6, 767–774.

Lill, N. L. & Sever, N. I. 2012. Where EGF receptors transmit their signals. Sci Signal, 5, pe41.

Maeda, K., Goto, S., Miura, K., Saito, K. & Morita, E. 2023. The Incorporation of Extracellular Vesicle Markers Varies Among Vesicles with Distinct Surface Charges. J Biochem.

Mathieu, M., Martin-Jaular, L., Lavieu, G. & Thery, C. 2019. Specificities of secretion and uptake of exosomes and other extracellular vesicles for cell-to-cell communication. Nat Cell Biol, 21, 9–17.

Mathieu, M., Nevo, N., Jouve, M., Valenzuela, J. I., Maurin, M., Verweij, F. J., Palmulli, R., Lankar, D., Dingli, F., Loew, D., Rubinstein, E., Boncompain, G., Perez, F. & Thery, C. 2021. Specificities of exosome versus small ectosome secretion revealed by live intracellular tracking of CD63 and CD9. Nat Commun, 12, 4389.

Matsui, T., Osaki, F., Hiragi, S., Sakamaki, Y. & Fukuda, M. 2021. Alix and ceramide differentially control polarized small extracellular vesicle release from epithelial cells. EMBO Rep, 22, e51475.

Miura, K., Jacques, K. M., Stauffer, S., Kubosaki, A., Zhu, K., Hirsch, D. S., Resau, J., Zheng, Y. & Randazzo, P. A. 2002. ARAP1: a point of convergence for Arf and Rho signaling. Mol Cell, 9, 109–19.

Moshiri, A., Humpal, D., Leonard, B. C., Imai, D. M., Tham, A., Bower, L., Clary, D., Glaser, T. M., Lloyd, K. C. & Murphy, C. J. 2017. Arap1 Deficiency Causes Photoreceptor Degeneration in Mice. Invest Ophthalmol Vis Sci, 58, 1709–1718.

Murayama, Y., Shinomura, Y., Oritani, K., Miyagawa, J., Yoshida, H., Nishida, M., Katsube, F., Shiraga, M., Miyazaki, T., Nakamoto, T., Tsutsui, S., Tamura, S., Higashiyama, S., Shimomura, I. & Hayashi, N. 2008. The tetraspanin CD9 modulates epidermal growth factor receptor signaling in cancer cells. J Cell Physiol, 216, 135–43.

Ostrowski, M., Carmo, N. B., Krumeich, S., Fanget, I., Raposo, G., Savina, A., Moita, C. F., Schauer, K., Hume, A. N., Freitas, R. P., Goud, B., Benaroch, P., Hacohen, N., Fukuda, M., Desnos, C., Seabra, M. C., Darchen, F., Amigorena, S., Moita, L. F. & Thery, C. 2010. Rab27a and Rab27b control different steps of the exosome secretion pathway. Nat Cell Biol, 12, 19–30; sup pp 1-13.

Pols, M. S. & Klumperman, J. 2009. Trafficking and function of the tetraspanin CD63. Exp Cell Res, 315, 1584–92.

Raposo, G. & Stoorvogel, W. 2013. Extracellular vesicles: exosomes, microvesicles, and friends. J Cell Biol, 200, 373–83.

Schoppe, J., Mari, M., Yavavli, E., Auffarth, K., Cabrera, M., Walter, S., Frohlich, F. & Ungermann, C. 2020. AP-3 vesicle uncoating occurs after HOPS-dependent vacuole tethering. EMBO J, 39, e105117.

Segeletz, S., Danglot, L., Galli, T. & Hoflack, B. 2018. ARAP1 Bridges Actin Dynamics and AP-3-Dependent Membrane Traffic in Bone-Digesting Osteoclasts. iScience, 6, 199–211.

Shiba, Y., Kametaka, S., Waguri, S., Presley, J. F. & Randazzo, P. A. 2013. ArfGAP3 regulates the transport of cation-independent mannose 6-phosphate receptor in the post-Golgi compartment. Curr Biol, 23, 1945–51.

Shiba, Y. & Randazzo, P. A. 2012. ArfGAP1 function in COPI mediated membrane traffic: currently debated models and comparison to other coat-binding ArfGAPs. Histol Histopathol, 27, 1143–1153.

Spang, A., Shiba, Y. & Randazzo, P. A. 2010. Arf GAPs: gatekeepers of vesicle generation. FEBS Lett, 584, 2646–2651.

Stenmark, H., Parton, R. G., Steele-Mortimer, O., Lutcke, A., Gruenberg, J. & Zerial, M. 1994. Inhibition of rab5 GTPase activity stimulates membrane fusion in endocytosis. EMBO J, 13, 1287–96.

Stuffers, S., Sem Wegner, C., Stenmark, H. & Brech, A. 2009. Multivesicular endosome biogenesis in the absence of ESCRTs. Traffic, 10, 925–37.

Su, M. Y., Fromm, S. A., Remis, J., Toso, D. B. & Hurley, J. H. 2021. Structural basis for the ARF GAP activity and specificity of the C9orf72 complex. Nat Commun, 12, 3786.

Sztul, E., Chen, P. W., Casanova, J. E., Cherfils, J., Dacks, J. B., Lambright, D. G., Lee, F. S., Randazzo, P. A., Santy, L. C., Schurmann, A., Wilhelmi, I., Yohe, M. E. & Kahn, R. A. 2019. ARF GTPases and their GEFs and GAPs: concepts and challenges. Mol Biol Cell, 30, 1249–1271.

Tomas, A., Futter, C. E. & Eden, E. R. 2014. EGF receptor trafficking: consequences for signaling and cancer. Trends Cell Biol, 24, 26–34.

Van Niel, G., Charrin, S., Simoes, S., Romao, M., Rochin, L., Saftig, P., Marks, M. S., Rubinstein, E. & Raposo, G. 2011. The tetraspanin CD63 regulates ESCRT-independent and -dependent endosomal sorting during melanogenesis. Dev Cell, 21, 708–21.

Venkateswarlu, K., Brandom, K. G. & Lawrence, J. L. 2004. Centaurin-alpha1 is an in vivo phosphatidylinositol 3,4,5-trisphosphate-dependent GTPase-activating protein for ARF6 that is involved in actin cytoskeleton organization. J Biol Chem, 279, 6205–8.

Verweij, F. J., Van Eijndhoven, M. A., Hopmans, E. S., Vendrig, T., Wurdinger, T., Cahir-Mcfarland, E., Kieff, E., Geerts, D., Van Der Kant, R., Neefjes, J., Middeldorp, J. M. & Pegtel, D. M. 2011. LMP1 association with CD63 in endosomes and secretion via exosomes limits constitutive NF-kappaB activation. EMBO J, 30, 2115–29.

Watanabe, A., Hataida, H., Inoue, N., Kamon, K., Baba, K., Sasaki, K., Kimura, R., Sasaki, H., Eura, Y., Ni, W. F., Shibasaki, Y., Waguri, S., Kokame, K. & Shiba, Y. 2021. Arf GTPase-activating proteins SMAP1 and AGFG2 regulate the size of Weibel-Palade bodies and exocytosis of von Willebrand factor. Biol Open, 10.

Wegner, C. S., Malerod, L., Pedersen, N. M., Progida, C., Bakke, O., Stenmark, H. & Brech, A. 2010. Ultrastructural characterization of giant endosomes induced by GTPase-deficient Rab5. Histochem Cell Biol, 133, 41–55.

Yoon, H. Y., Kales, S. C., Luo, R., Lipkowitz, S. & Randazzo, P. A. 2011. ARAP1 association with CIN85 affects epidermal growth factor receptor endocytic trafficking. Biol Cell, 103, 171–84.

Yoon, H. Y., Lee, J. S. & Randazzo, P. A. 2008. ARAP1 regulates endocytosis of EGFR. Traffic, 9, 2236–52.

